# Distinct cortical correlation structures of fractal and oscillatory neuronal activity

**DOI:** 10.1101/2020.12.10.415315

**Authors:** Andrea Ibarra Chaoul, Markus Siegel

## Abstract

Electrophysiological population signals contain oscillatory and fractal (1/frequency) components. So far research has largely focused on oscillatory activity and only recently interest in fractal population activity has gained momentum. Accordingly, while the cortical correlation structure of oscillatory population activity has been characterized, little is known about the correlation of fractal neuronal activity. To address this, we investigated fractal neuronal population activity in the human brain using resting-state magnetoencephalography (MEG). We combined source-analysis, signal orthogonalization and irregular-resampling auto-spectral analysis (IRASA) to systematically characterize the cortical distribution and correlation of fractal neuronal activity. We found that fractal population activity is robustly correlated across the cortex and that this correlation is spatially well structured. Furthermore, we found that the cortical correlation structure of fractal activity is similar but distinct from the correlation structure of oscillatory neuronal activity. Anterior cortical regions showed the strongest differences between oscillatory and fractal correlation patterns. Our results suggest that correlations of fractal population activity serve as robust markers of cortical network interactions. Furthermore, our results show that fractal and oscillatory signal components provide non-redundant information about large-scale neuronal correlations. This may reflect at least partly distinct neuronal mechanisms underlying and reflected by oscillatory and fractal neuronal population activity.

## Introduction

Neuronal population activity as measured with MEG, EEG or local field potentials (LFP) consists of oscillatory and fractal signal components. These components are evident from the power spectra of such signals, which show a characteristic 1/frequency shape, corresponding to fractal signal components, and variable peaks of oscillatory activity.

Cortical oscillations have attracted strong interest and have been implicated in various functions [1–5]. The amplitudes of neuronal oscillations show characteristic patterns of correlation across the cortex [6–9]. These patterns are similar to BOLD fMRI correlation patterns [10], are distinct from phase-coupling patterns [8], and serve as biomarkers of various neuropsychiatric diseases, such as e.g. major depressive disorder [11], schizophrenia [12–14], autism [15], Alzheimer’s disease [16] and congenital blindness [17].

In contrast, little is known about fractal neuronal population activity. It is not until recently that studies of the biological importance of fractal activity gained momentum [18]. Intra-cortically, fractal power is closely linked to neuronal spiking [19,20] and allows to decode motor behavior and visual inputs with even better fidelity than neuronal oscillations [21–23]. Non-invasively, EEG and MEG studies have shown that the slope of the fractal component is modulated by sleep [24], drugs [25,26], ageing [27], anesthesia [28,29], and meditation [30]. However, in contrast to oscillatory activity, nothing is known about the cortical coupling of the amplitude of fractal population activity.

To address this, we investigated two specific questions. First, what is the cortical correlation structure of fractal activity in the human brain? And second, how does the correlation structure of fractal activity compare to that of oscillatory population activity?

To address these questions, we combined human resting state MEG, source reconstruction, signal orthogonalization and analytical techniques for separating oscillatory and fractal components. We found that spontaneous co-fluctuations of cortical fractal activity in the human brain are spatially well structured and that, although similar, there are consistent differences between the correlation structure of fractal and oscillatory neuronal population activity. Our results suggest correlations of fractal population activity as robust markers of large-scale cortical interactions.

## Results

We analyzed combined data from two datasets of resting state MEG recordings in healthy human participants (n = 112). One dataset (n = 89 subjects) was from the Human Connectome Project [31,32] and the other dataset (n = 23) was recorded at the MEG Center, Tübingen.

### Fractal cortical population activity

We source reconstructed cortical activity from the MEG data using beamforming [33] and separated oscillatory and fractal components using irregular-resampling auto-spectral analysis (IRASA) [34]. IRASA extracts the scale free characteristic, i.e. spectral self-similarity, of the broadband signal. It applies a series of re-samplings to generate power spectra that displace the oscillatory peaks, but do not alter the scale free aspect of the spectrum. The fractal component is then derived as the median over all re-samplings (see Methods). IRASA has several key advantages. It is non-parametric, i.e. there is no need to pre-specify the number or width of oscillatory modes, it can automatically detect different fractal regimes and it operates across a large frequency range.

Fractal power of human brain activity has been described either by a single power law or by a model with two power laws joined with a knee [18,23,24,25,34,35,51]. Importantly, MEG recordings also pick-up ambient noise, which can be power law shaped [36] and may thus affect estimates of fractal population activity. Thus, before investigating the co-fluctuation of fractal activity, we first aimed to dissociate these factors by modelling the fractal power spectrum with a parametric model. Furthermore, we determined the number of fractal regimens using model selection, which has biological implications that may help to elucidate the mechanisms underlying the fractal power spectrum. We fitted three increasingly complex models: a single power law (Fig 1B, top), two power laws connected by a knee at 15 Hz (Fig 1B, bottom), and two connected power laws with added noise of empty room MEG recordings (Fig 1A).

**Fig 1.**
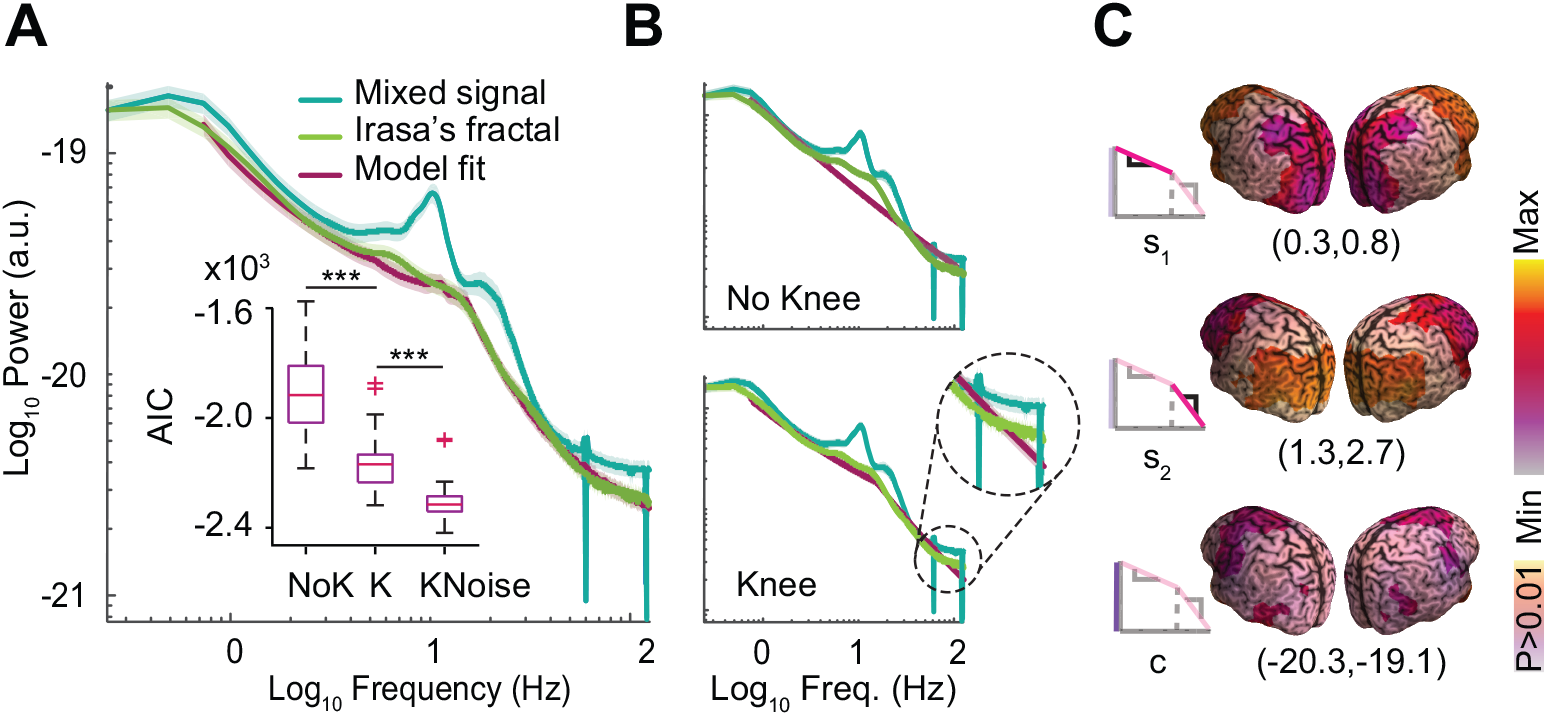
Fractal cortical population activity. (**A**) Average power spectrum of the mixed broad-band signal, of the fractal signal component isolated by IRASA and of the optimal model of fractal power (two power laws with knee and noise). The inset shows the AIC for the three different models (all comparisons P < 10^−14^, n = 109, two tailed t-test). (**B**) Model fits for one power law (no knee) and two power laws (knee) fail to capture the shape of the fractal power spectrum. Shaded error bars denote SEM across subjects. (**C**) Cortical distribution of the three neuronal parameters (*s_1_*, *s_2_* and *c*) of the optimum model (see Fig S1 for noise coefficient). *s_1_*,: 0.7 - 15 Hz slope, *s_2_*: 15 - 125 Hz slope, *c*: constant term. P-values are from cluster permutation statistics across subjects (n = 109, 1000 permutations).

The third model with two connected power laws and noise best described the data and was optimal according to the Akaike Information Criterion (AIC) (Fig 1A, inset) (AIC one power law vs. AIC two power laws: P < 10^−15^, t-test, n = 109; AIC two power-laws vs AIC two power-laws and noise P < 10^−15^, two tailed t-test, n = 109). This optimal model had four parameters (Fig 1C): the two slopes of the power laws (s_1_ and s_2_), the offset of the power (c), and the amount of MEG noise (see Fig S1A for the cortical distribution of the noise component).

The model revealed a highly specific cortical distribution of fractal neuronal activity (Fig 1C). All four model parameters had consistent non-random cortical patterns across subjects (all P < 0.01, cluster permutation test). The 0.7 to 15 Hz power-law slope was steepest in anterior areas with minima in sensorimotor and visual cortex (s_1_; 0.46 +/− 0.12, mean +/− SD). In contrast, the 15 to 125 Hz slope was steepest in posterior regions, excluding the occipital pole (s_2_; 1.99 +/− 0.39, mean +/− SD). The constant term (c; −19.6, +/− 0.3, mean +/− SD) was lowest in prefrontal and medial temporal cortex and the noise term (n; 0.5 +/− 0.09, mean +/− SD) peaked in the temporal regions that are often contaminated by muscle artifacts (Fig S1A).

The present MEG data consisted of two independent datasets recorded with different MEG systems at different sites. Thus, we next tested if results were consistent across datasets. This was indeed the case. For both datasets independently, the two power-law model including the noise component was the optimal fractal model (all model comparisons P < 10^−4^, two tailed t-test, n=109). Also, the specific model parameters were consistent across datasets. In fact, the first and second slope of the fractal model as well as the noise coefficient were not significantly different between the two datasets (all P > 0.05, t-tests). Only the constant term showed a significant difference (P = 2×10^−7^, t-test; HCP: −19.45+/−0.18, Tübingen: −20.04+/−0.17).

In summary, consistent across two independent MEG datasets we identified fractal neuronal population activity that was well described by a two power-law model. The spectral shape and strength of this fractal activity showed a specific cortical distribution with anterior-posterior gradients.

### Oscillatory cortical population activity

If IRASA performed a valid separation of broad-band signal power, the retained oscillatory components should resemble the known spectral and cortical specificity of rhythmic brain activity. Indeed, this is what we found. Fig 2A shows the oscillatory power for three characteristic seeds: left auditory, left somatosensory and medial prefrontal cortex with peaks in the alpha, beta and theta band, respectively. The cortical distribution of oscillatory power (Fig 2B) showed strongest theta band activity (6 Hz) in frontal areas, a prominent alpha peak (11.3 Hz) over visual cortex and a beta peak (22.3 Hz) over sensorimotor cortices (all P < 0.05; cluster permutation test). Thus, oscillatory power derived with IRASA well resembled the expected spectral and cortical profile of cortical rhythms.

**Fig 2.**
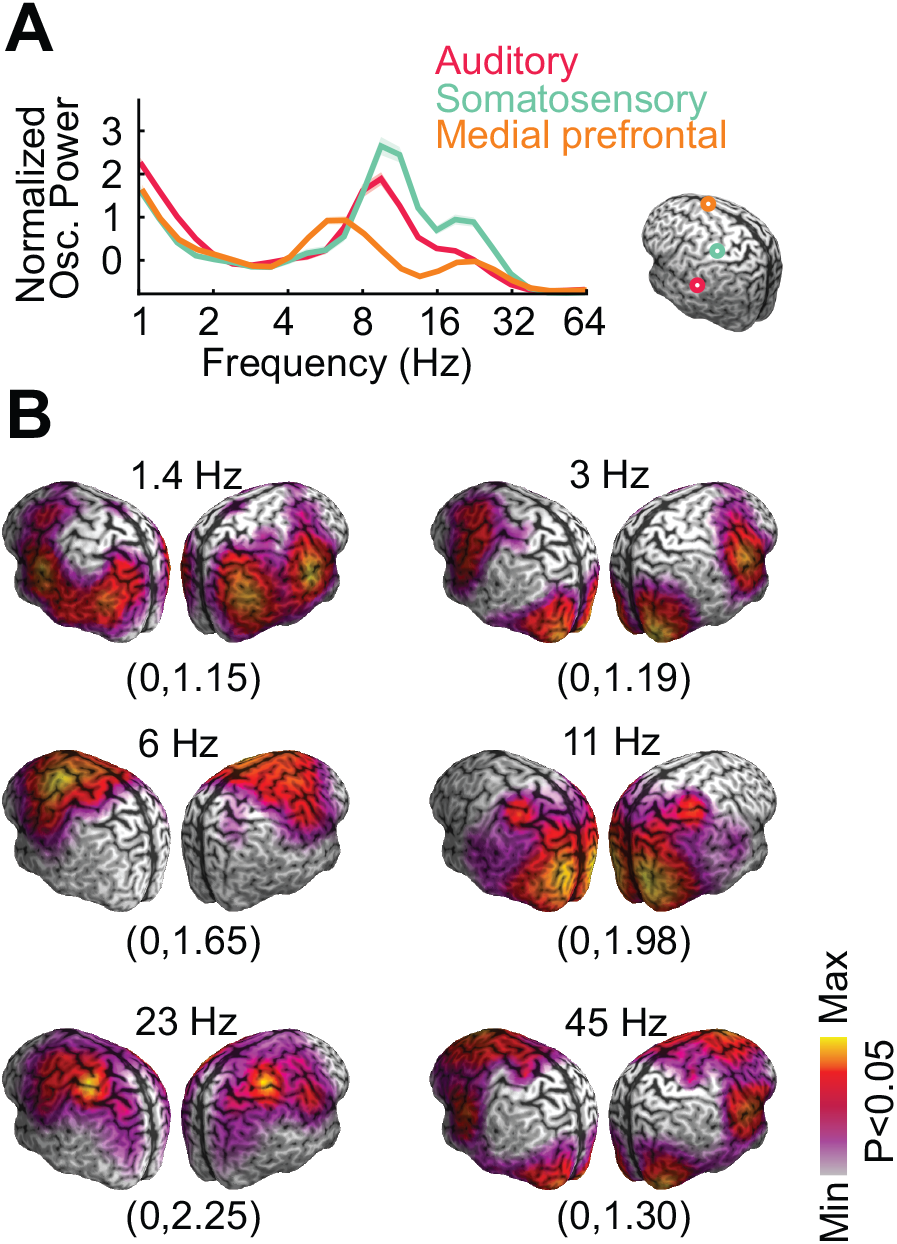
Oscillatory cortical population activity. (**A**) Power spectra of the oscillatory component derived by IRASA for left auditory cortex, left somatosensory cortex and medial prefrontal cortex. (**B**) Cortical distribution of oscillatory power. P-values are from cluster permutation tests across subjects (n=109).

### Cortical correlation structure of fractal activity

We next turned to our first question – if the correlation of fractal cortical activity is spatially structured. Signal leakage confounds the direct correlation of MEG signals. Thus, we applied pairwise orthogonalization of source-reconstructed MEG signals to rule out such confounds [6–8]. We confirmed that orthogonalization suppressed leakage effects (Fig S2) but did not alter the power spectrum (Fig S3). We then applied IRASA across time using a sliding window approach (3 s windows), fitted the fractal model for each time window and analyzed cortical co-fluctuations of activity across time for the different parameters of the fractal model.

For each cortical location and fractal model parameter, we quantified its mean correlation to all other cortical locations (Fig 3A). We found that all fractal parameters had a consistent cortical distribution across subjects and peaked in parietal cortex (P < 0.01, cluster permutation test; see Fig S1 for noise component). Thus, overall parietal cortex showed the strongest correlation of fractal activity to other brain regions and may thus function as a hub of fractal power correlations. While overall the correlation structure was similar for all model parameters, descriptively s_1_ peaked more anterior than s_2_, which further supports the need to employ a two power-law model.

**Fig 3.**
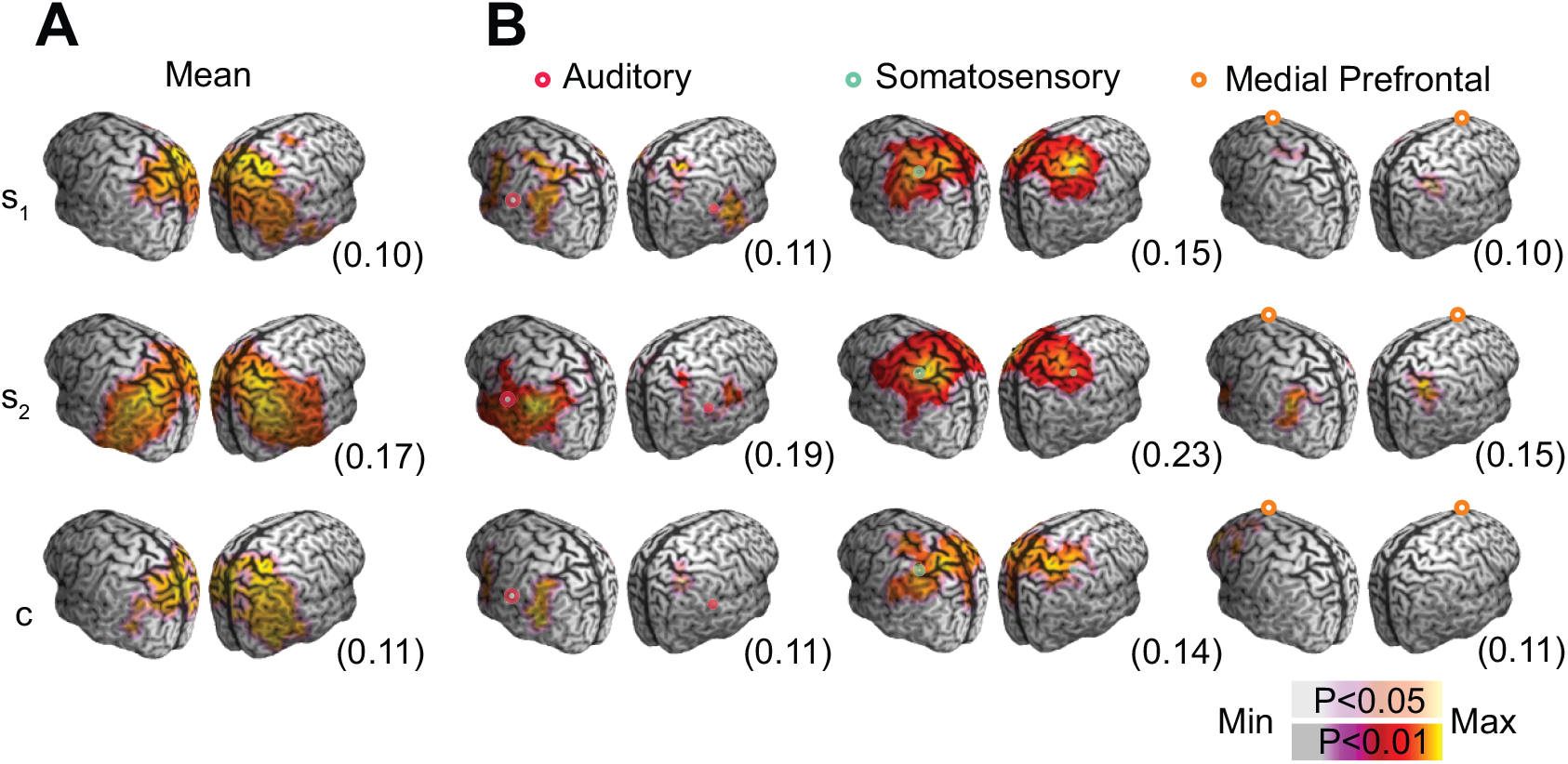
Cortical correlation structure of fractal activity. (**A**) Cortical distribution of the average correlation of fractal model parameters across time (*s_1_*: 0.7-15 Hz slope, *s_2_*: 15-125 Hz slope, *c* constant term). Maximum correlations are in parentheses. (**B**) Cortical correlation structure of fractal model parameters for three cortical seeds (left auditory cortex, left somatosensory cortex and medial prefrontal cortex). In parentheses are the maximum correlations values. All P-value are from cluster permutation tests, masked at P < 0.05, n=107.

We next analyzed the cortical correlation pattern of fractal parameters for specific cortical regions (Fig 3B). We focused on early auditory and somatosensory cortex, which show strong interhemispheric coupling for fMRI [37], intracranial recordings [38] and rhythmic M/EEG activity [6,9]. Furthermore, we investigated medial prefrontal cortex, which shows rhythmic coupling with frontoparietal cortices [6,8]. If fractal signal components are involved in functional interactions between brain regions, these components should show similar correlation structures. This is what we found (Fig 3B). Across model parameters, all seeds showed the expected bilateral correlations structures with most distinct patterns for somatosensory cortex. Furthermore, correlation structures were generally consistent across model parameters with strongest correlation patterns for the 15 to 125 Hz slope (s_2_). In summary, we found that spontaneous fluctuations of fractal power were correlated across the brain. These correlations were spatially well structured, with parietal regions as major hubs and with prominent interhemispheric correlations of homologous sensory areas.

### Comparison of fractal and oscillatory correlation structures

After establishing that the cortical correlation of fractal activity was indeed spatially structured, we focused on our second question – how does the correlation structure of fractal activity compare to the correlation structure of oscillatory activity? In order to perform this comparison in a frequency-specific fashion, we reconstructed the power spectrum of fractal activity from the fractal model, discarding the noise component (Fig 4). In analogy to the fractal model parameters above, we then correlated the power of the fractal and oscillatory component between all pairs of brain regions across time. In addition, we performed the same analysis for the original mixed signal, that contained both, fractal and oscillatory components.

**Fig 4.**
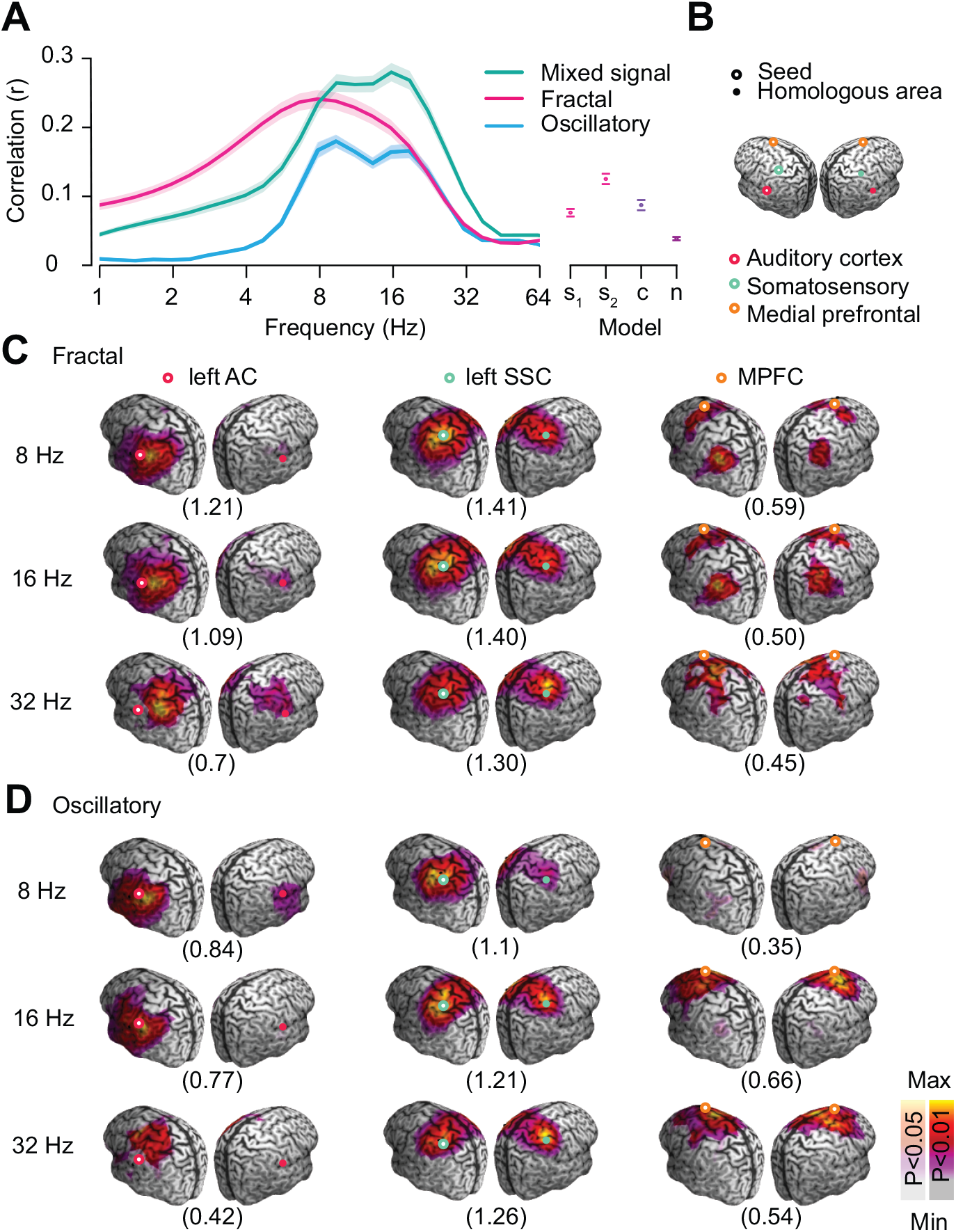
Comparison of fractal and oscillatory correlation patterns. (**A**) Average correlation of the mixed cortical signal as well as of oscillatory and fractal signal component. The mean correlations of the individual fractal model parameters are on the right. Shaded areas and error bars denote SEM across subjects. (**B**) Cortical seed locations. (**C**) Cortical correlation structure of fractal activity for three seeds and frequencies. (**D**) Cortical correlation structure of oscillatory activity. Numbers in parenthesis are the maximum t-scores of significant clusters. All P-values are from cluster permutation tests, n = 107.

Averaged across all pairs of brain regions, oscillatory activity showed a spectrally narrower and bimodal distribution of correlation than fractal activity (Fig 4A). However, correlations were generally stronger for fractal signal components (see Fig S4 for correlations of the raw fractal output of IRASA and of the suboptimal fractal models).

Next, we directly compared the cortical correlation patterns between fractal and oscillatory activity for the same seed regions that we employed for the fractal model parameters for three frequencies with strong average correlations (Fig 4C and D). In accordance with the correlation structure of the fractal model parameters, we found strong interhemispheric correlations for auditory and somatosensory cortex as well as bilateral frontoparietal correlations for medial prefrontal cortex. In general, these correlation structures were similar between fractal and oscillatory activity. However, fractal correlations were stronger and with more distinct spatial structure. Thus, we next systematically quantified the similarity of fractal and oscillatory correlation structures (Fig 5).

**Fig 5.**
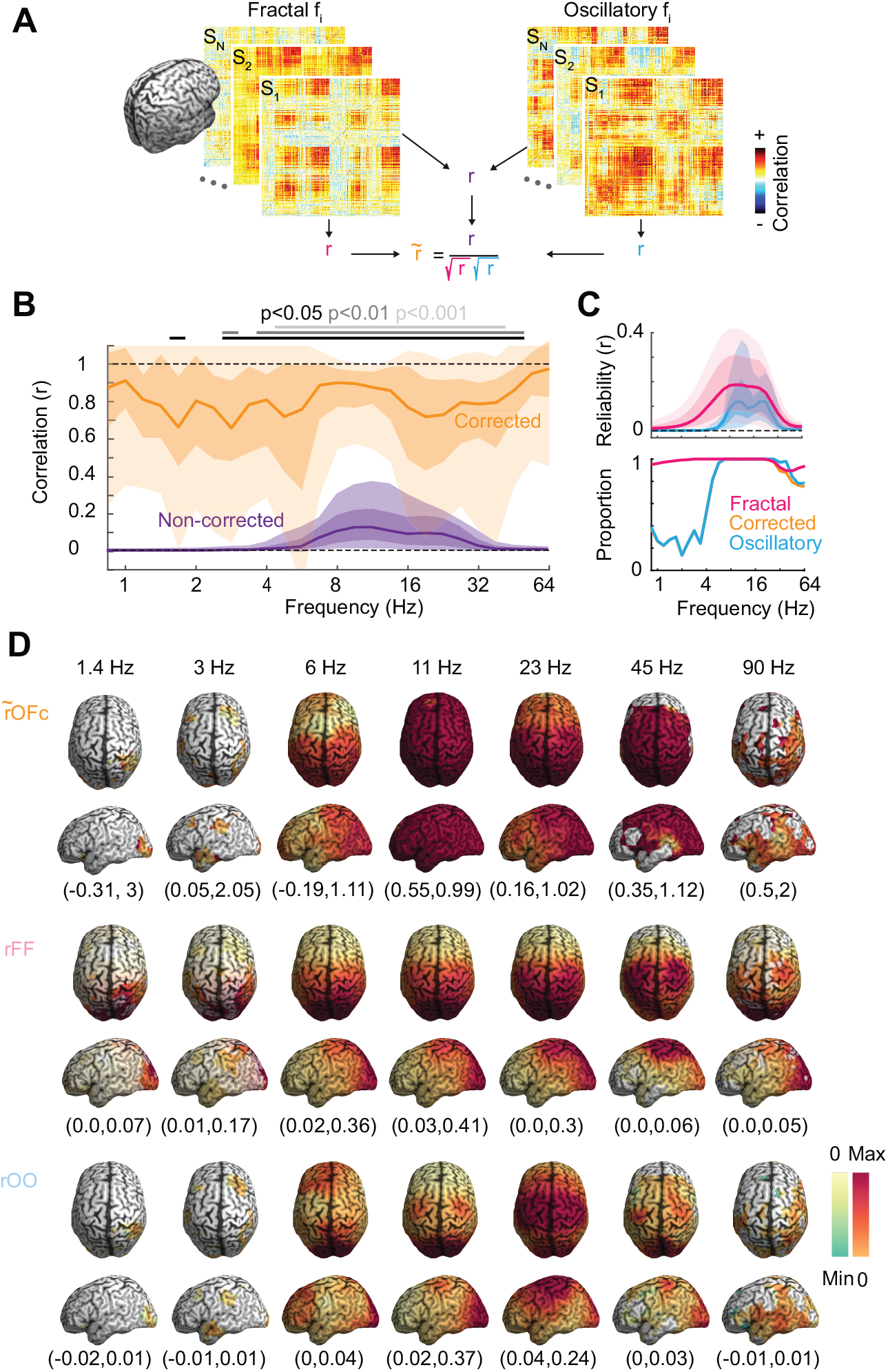
Attenuation corrected correlation of fractal and oscillatory correlation patterns. (**A**) Diagram of attenuation correction procedure. (**B**) Corrected and non-corrected median correlation. Shaded regions denote 5-95^th^ and 25-75^th^ percentiles across the cortex. Significance bars indicate corrected correlations < 1 (FDR corrected, n=107, one tailed t-test). (**C**) Fractal and oscillatory reliabilities (top), and the proportion of cortical seeds with reliable correlation patterns (bottom) (**D**) Cortical distribution of attenuation corrected correlation between oscillatory and fractal correlation (rOFc) and reliabilities (rFF for fractal and rOO for oscillatory for different frequencies. Minimum and maximum corrected correlations are in parentheses. Desaturated or gray regions have <50% or no reliable subjects, respectively.

### Fractal and oscillatory correlation patterns are distinct

For each seed region and frequency, we correlated the correlation patterns of fractal and oscillatory components (Fig 5B, purple). Averaged across all seed regions, correlation coefficients were positive and up to 0.2. Furthermore, correlation patterns were most similar in the frequency range between 8 and 16 Hz. This may indicate that oscillatory and fractal correlations pattern are most similar in this frequency ranges. However, the measured correlation between two quantities does not only reflect their underlying true correlation but also the reliability (or signal-to-noise ratio) with which these quantities are measured. Thus alternatively, the peaked correlation between 8 and 16 Hz may merely reflect the higher reliability of correlation patterns in this frequency range.

We applied attenuation correction of correlations [39] to disambiguate these alternatives and to discount the effect of reliability (Fig 5A; for other recent uses of this approach see: [8,9,40]. To enable population-level inferences, we estimated the reliability of fractal and oscillatory correlation patterns as their inter-subject correlation. High reliability indicates consistent correlation patterns across subjects, whereas low reliability indicates strong variability across subjects. Fractal and oscillatory reliability both peaked from about 8 to 16 Hz with stronger reliability for fractal correlation patterns and two peaks for oscillatory reliability around 10 and 23 Hz (Fig 5C). Fractal reliability was broadly distributed across the cortex peaking in parietal, occipital and central regions (Fig 5D). Oscillatory reliability showed a more distinct pattern with frequency-dependent peaks in frontal, central and occipital regions.

We used these reliabilities to correct the correlation between oscillatory and fractal correlation patterns (Fig 5A). This had a marked effect. The corrected correlation coefficient between oscillatory and fractal correlation patterns was around 0.8 with little variability across frequencies (Fig 5B; orange). This had two implications. First, the spectral specificity of the similarity between oscillatory and fractal correlation patterns was largely driven by different reliabilities across frequencies. Second, the fact that corrected correlation coefficients were high, but significantly smaller than 1 across a broad frequency range (P < 0.01 corrected, one tailed t-test, n = 107; Fig 5B), implied that cortical correlation patterns of oscillatory and fractal activity were similar but distinct.

### Differences between fractal and oscillatory correlation patterns point to known oscillations

Given that fractal and oscillatory correlation patterns are distinct, how can we characterize their difference? We addressed this question in two ways. First, we compared fractal and oscillatory correlation patterns across frequencies, and second, at the connection level.

In the first approach, i.e. the comparison across frequencies, we first isolated the differences between fractal and oscillatory correlation patterns and then compared these differences across frequencies (Fig 6). For each frequency and subject, we normalized and subtracted fractal and oscillatory correlation patterns, which removed their global similarities (see also Fig S5A). We then correlated the differences between fractal and oscillatory correlation patterns across frequencies (Fig 6A). In other words, we quantified if what made fractal and oscillatory correlation patterns different at one frequency, was similar to the difference at another frequency. Importantly, we again corrected for reliability confounds by applying attenuation correction (see also Fig S5B).

**Fig 6.**
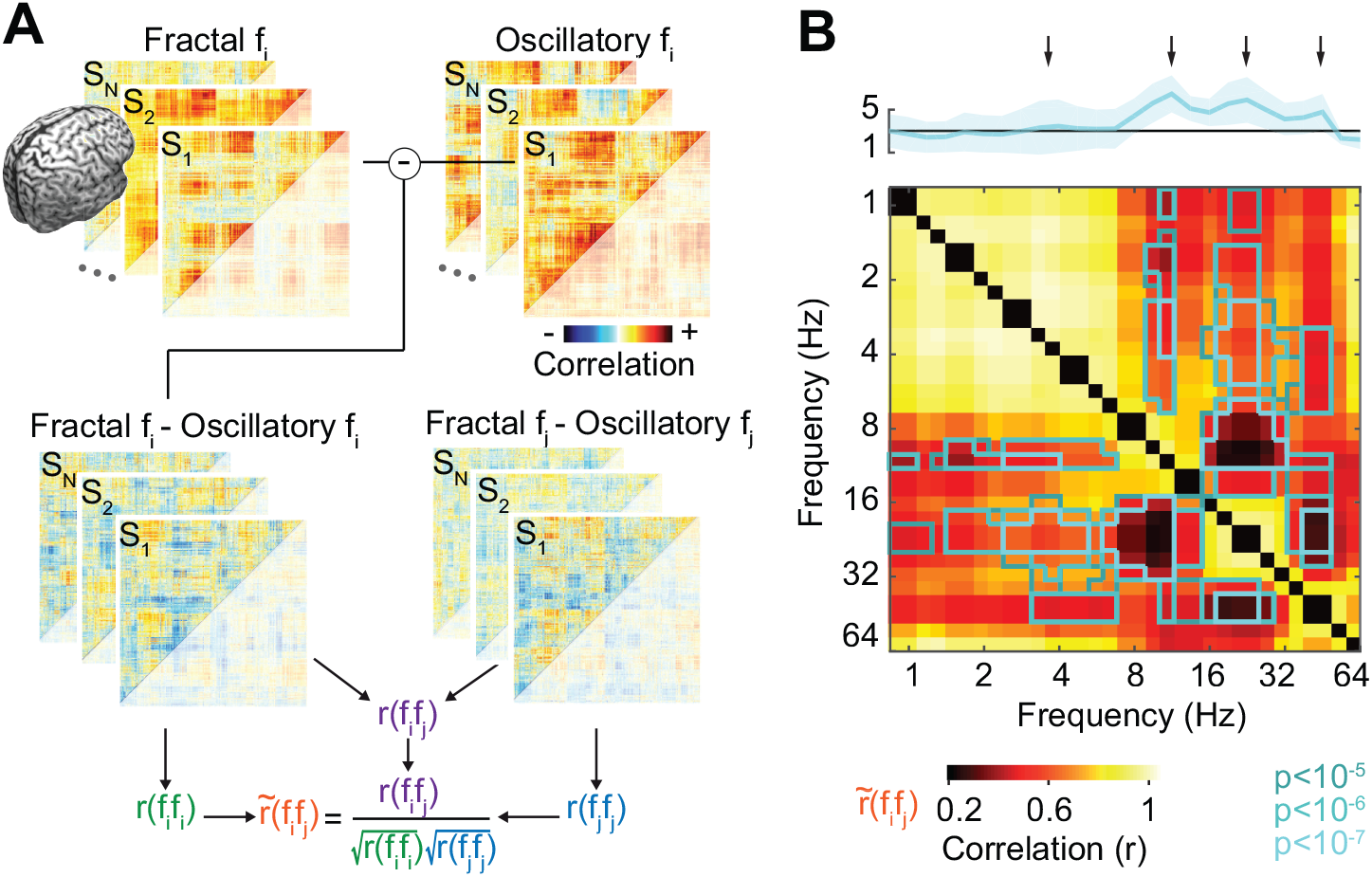
Correlation of pattern differences across frequencies. (**A**) Diagram of the attenuation correction procedure applied to the differences of connectivity matrices of fractal and oscillatory components across frequencies. (**B**) Corrected cross-frequency correlation of differences between fractal and oscillatory correlation patterns. Outlined areas show corrected correlation values <1 (FDR corrected, n = 107, one tailed t-test). The cyan spectrum at the top is the average negative log10 of the p-values per frequency. The arrows indicate the natural spectral grouping of differences.

We found that, across the entire frequency space, corrected correlation coefficients were broadly distributed between 0.2 to 1 (Fig 6B). Attenuation correction allowed us to test, which frequency combinations showed distinct differences between fractal and oscillatory correlations, i.e. correlation coefficients smaller than 1. If organized in columns, we can think of these frequencies as separators of distinct differences between fractal and oscillatory correlations. Indeed, testing for correlations smaller than 1 (P < 10^−5^ FDR corrected, one tailed t-test, n = 107) revealed a clear columnar structure separating frequency ranges with known brain rhythms. For example, 10 Hz showed correlations smaller than 1 with all lower frequencies (~0.8 to ~6.7 Hz) and frequencies higher than 16 Hz. This identifies 10 Hz as having consistent differences between oscillatory and fractal correlations that only occur at 10 Hz and neighboring frequencies. We found similar separations for frequencies around 4 Hz, 20 Hz and 40 Hz. Differences around 4 Hz were distinct from 10 to 45 Hz but not from other frequency ranges.

In sum, the comparison across frequencies identified cortical delta, alpha, beta and low gamma frequency bands and showed that the correlation structure of oscillatory activity in these bands had distinct differences to the correlation of fractal neuronal activity.

### Local differences between fractal and oscillatory correlations

We next compared fractal and oscillatory correlations at the connection level to identify which specific connections were distinct. For this, we employed local attenuation corrected correlation [40]. In this approach, not the entire cortex-wide correlation patterns were correlated between activities, but instead we analyzed for each connection, i.e. each pair of regions, only the correlation patterns of these regions’ local neighborhoods. E.g., figure 7A shows the neighborhoods surrounding seeds in left prefrontal and right parietal cortex. This approach discounts coarse global correlation patterns and allows to resolve the similarity of correlation patterns between two specific regions.

**Fig 7.**
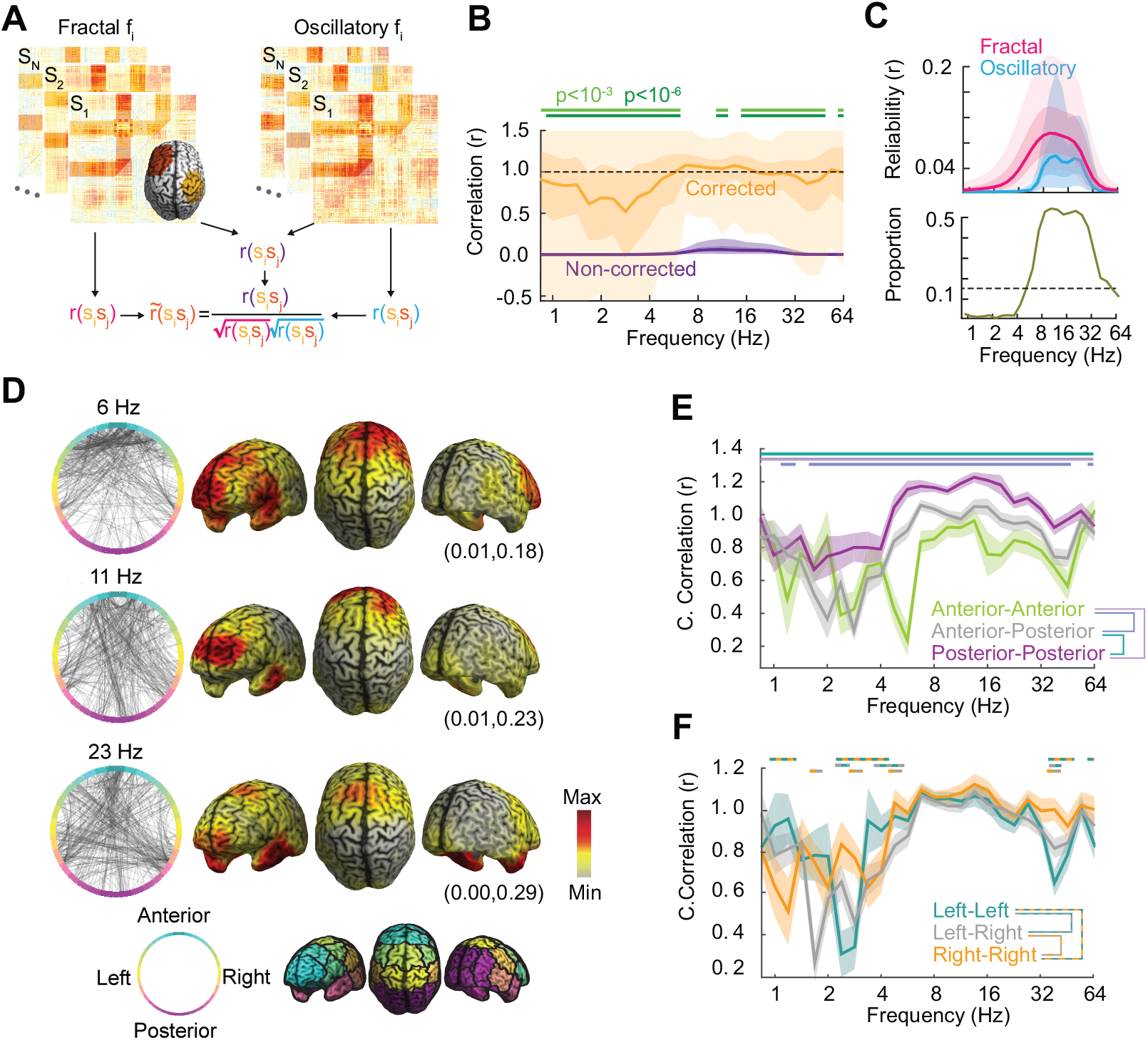
Differences of correlation at the connection level. (**A**) Schema of local attenuation correction showing the neighborhood of two example seeds. (**B**) Median corrected and non-corrected correlation of local oscillatory and fractal correlation patterns. Shaded regions show 9-95^th^ and 25-75^th^ percentiles. P-values are for tests < 1 (FDR-corrected, n = 107, one tailed t-test). (**C**) Average reliabilities and proportion of reliable connections of oscillatory and fractal correlations. (**D**) Circle plots show a random sample (800 connections) of the 10 % lowest average corrected correlations across frequency-bands and subjects. The topographies show the marginal mean of the corrected correlations that are in the 10 % lowest correlations per frequency band. Minimum and maximum marginal correlations are in parentheses. (**E**) Mean corrected correlation of local oscillatory and fractal correlation patterns grouped into anterior-anterior, posterior-posterior and anterior-posterior connections. Bars indicate pair-wise differences (P < 0,05 FDR corrected; shaded areas denote SEM across subjects, n = 107, one tailed t-test). (**F**) Same as (E) but for connections separated into left-left, left-right and right-right groups.

In accordance with the global correlation analysis, local attenuation corrected correlation showed similar fractal and oscillatory correlation patterns that were however significantly distinct (correlation < 1) across broad frequency ranges (Fig 7B; P < 10^−3^ and p<10^−6^ corrected, one tailed t-test, n=107). Patterns were most similar in the alpha frequency range around 10 Hz, where the average correlation was indeed not significantly smaller than 1. Again, local reliabilities peaked from about 8 to 16 Hz and were stronger for fractal than for oscillatory correlations (Fig 7C).

To answer which connections underlie the differences between fractal and oscillatory correlations, we identified from all the reliable connections, the 10 % least similar connections for each frequency (Fig 7D). From 6 to 23 Hz, where at least 15 % of connections were reliable, dissimilar connections were mostly within frontal and between frontal and temporal cortices (see Fig S6 for all individual frequencies and the average across all frequencies). This suggested that specifically anterior cortical regions showed distinct fractal and oscillatory correlations patterns.

To quantitively assess this, we divided cortical connections into three groups: anterior-anterior, posterior-posterior and anterior-posterior. We then compared the correlation between fractal and oscillatory correlation patterns between these three groups (Fig 7E). Indeed, we found that anterior-anterior connections were most dissimilar, followed by anterior-posterior connections, and posterior-posterior connections. Furthermore, this analysis revealed a marked dissimilarity between fractal and oscillatory correlations specifically for anterior-anterior connections around 6 Hz. A control analysis separating connections according to left and right hemisphere (Fig 7F) did not reveal a similar pattern.

In sum, the comparison on the connection level identified cortical correlations involving the frontal cortex as most dissimilar between fractal and oscillatory neuronal activity.

## Discussion

We combined source-analysis, signal orthogonalization and IRASA to systematically characterized the cortical distribution and correlation of fractal neuronal activity in resting state MEG. We found that fractal population activity is robustly correlated across the cortex, that this correlation is spatially well structured, and that the cortical correlation structure of fractal activity is similar but distinct from the correlation structure of oscillatory population activity.

### Spectral profile of fractal activity

Our findings add to previous studies that have characterized the spectral profile of fractal cortical activity as multi-fractal [23–25,41]. Based on objective criteria (AIC) we identified an optimal model consisting of two continuous power laws with a knee at 15 Hz and noise. The continuous piece-wise linear model reduced the number of parameters and allowed to effectively reconstruct the fractal power spectrum with one constant term and two slopes. Furthermore, modelling the noise [23,36] did not only critically improve the model fit, but also allowed to reconstruct noise-free fractal power, which added an additional degree of independence between oscillatory and fractal activity.

The average slope coefficients that we identified were generally more extreme than those previously reported for MEG. Specifically, the average low- and high-frequency slopes (<15 Hz: 0.46; >15 Hz: 1.99) were smaller and larger than previously reported, respectively [25,34,42]. This could be explained by several factors: First, in contrast to most previous studies, we separated oscillatory and fractal components before quantifying fractal slopes. Second, the exact frequency range over which slopes are estimated differs between studies. Third, in contrast to previous studies [25] we measured resting state with eyes open. Fourth, in contrast to previous studies [24,25] we modeled the noise component of the power spectrum. In particular at very low and high frequencies, modeling the noise component assigns power to the noise term. This leads to smaller and larger fractal slopes for low and high frequencies, respectively.

Intriguingly, the spectral profile of fractal power that we found is closer to the profile that has been reported for intra-cortical measurements, which are less affected by noise [23,25]. Thus, our approach which includes component separation (IRASA) and noise modelling may yield measurements of fractal activity closer to the underlying intracortical activity.

### Cortical profile of fractal activity

In accordance with previous reports [25,34,43,50,51], source analysis and parametric modelling of fractal activity showed a robust and prominent anterior-posterior gradient of the cortical distribution of the fractal slopes. In particular, we found steeper fractal slopes for frequencies below and above 15 Hz in anterior and posterior regions, respectively. Further large-scale invasive studies are required to link these findings to measures of local populations activity (LFP) and spiking activity.

### Cortical correlation structure of fractal activity

This study provides, to our knowledge, the first systematic characterization of the cortical correlation structure of fractal activity. Our results extend previous reports that linked correlations of fractal activity to fMRI signals [24]. Importantly, we combined source reconstruction (beamforming) and signal orthogonalization to discount spurious correlations generated by signal leakage and to directly characterize the correlation structure at the cortical level. We found that temporal fluctuations of fractal activity were robustly correlated across the cortex and this these correlations were spatially well structured. Average correlations were strongest in parietal cortex with prominent interhemispheric correlations of homologous sensory areas and frontoparietal cortices. These results suggest that correlations of fractal population activity serve as robust markers of large-scale cortical network interactions.

### Comparison of fractal and oscillatory component correlations

Our results provide a systematic comparison of the correlation structure of fractal and oscillatory activity across the cortex and a broad frequency range. This extends previous studies reporting correlations between oscillatory and fractal components mostly in the alpha band [24,25]. Importantly, we employed attenuation correction of correlations to quantify unbiased correlations independent of reliability confounds. On the one hand, this approach revealed a high similarity of fractal and oscillatory correlation structures. On the other hand, this approach allowed to identify significant differences between both correlation structures.

We characterized these differences in two ways. First, we performed a cross frequency attenuation correction of the differences of oscillatory and fractal correlation. We found differences of correlation structures at known oscillatory frequencies. This accords well with the spectral specificity of unseparated, broad-band power correlations [6]. Furthermore, it provides evidence for a successful signal separation because fractal correlation patterns are expected to generalize across the frequency ranges of known cortical oscillations.

Second, we compared local fractal and oscillatory correlation patterns. This revealed that the both, the similarity and differences between both patterns were also present at the local connection level and did thus not merely reflect global gradients or trends. This analysis yielded two further insights. First, fractal and oscillatory correlations are most dissimilar for anterior cortex in the theta band (around 6 Hz). Second, and in contrast, correlations of alpha-band activity within posterior cortex were not different between fractal and oscillatory activity. This suggests that there is no generic link between co-fluctuation of oscillatory and fractal neuronal activity, but that this relationship is specific for the cortical rhythm and region at hand.

### Relationship between fractal activity and alpha oscillations

Throughout our analyses, we noted close links between fractal and oscillatory activity in the alpha band (around 12 Hz). The attenuated corrected correlation between oscillatory and fractal correlation patterns peaked around 12 Hz (Fig 5B). In the cross-frequency analysis of pattern differences (Fig S5C), reliabilities around 12 Hz decreased despite strong reliabilities within each signal component (Fig 5C). Finally, in the local analysis, differences in the alpha band were not over visual or motor cortices (Fig 7), which display strong alpha-band activity. Instead, the correlation patterns of posterior alpha-band activity and fractal activity were not different.

Several factors may explain these links. First, IRASA may not entirely separate oscillatory and fractal components which may drive correlations between both components. Second, the fractal part of the M/EEG spectrum may at least partially reflect the damped nature of alpha oscillators, which may cause correlations between alpha-band power and reconstructed fractal activity [25]. Third, spectral leakage of alpha oscillations into neighboring frequencies may result in correlations of oscillatory and fractal components.

### Similarity of fractal and oscillatory correlation patterns

Our results show strong similarities between the correlation structures of oscillatory and fractal cortical population activity as measured with MEG. This is in contrast to results from intracortical recordings [44,45], which suggest a stronger dissociation between oscillatory and fractal components. The different signal modalities may explain this discrepancy.

First, MEG may only weakly reflect the underlying fractal neuronal activity. For ECoG, it has been suggested that, compared to oscillatory spiking, fractal spiking has a smaller influence on the population signal [44], and that this difference is further accentuated with distance to electrode. Thus, the fractal signal picked up by MEG sensors may only to a small extent reflect to underlying fractal spiking activity. Second, along a similar line, fractal activity may indeed be correlated with oscillatory activity, but at a spatial scale coarser than assessed with LFP or ECoG recordings. MEG may be particularly sensitive to these coarser correlations. Finally, the employed behavioral task may have an important effect. We studied resting state activity, while invasive studies focused on correlates of motor behavior [45] and sensory stimulation [44]. Visual stimulation can robustly drive high-frequency MEG activity above 100 Hz [46,47], which may reflect broad-band spiking that is in turn dissociated from oscillatory activity.

### Differences of fractal and oscillatory correlation patterns

We found that oscillatory correlation patterns were dissimilar across broad frequency ranges on the global and local level. Independent of the exact mechanisms underlying the fractal part of the M/EEG signal, these differences may reflect distinct neurophysiological mechanisms underlying both signal components. Moreover, these findings imply that fractal and oscillatory signal components contain non-redundant information. Thus, the separation of power into its fractal and oscillatory components might be useful step for the study of normal and diseased brain function. Correlations of oscillatory activity serve as biomarkers for various neuropsychiatric diseases [11][12–14][15][17]. Correlations of fractal activity may allow to further improve these biomarkers.

## Methods

### MEG datasets

We analyzed data from 112 healthy subjects recorded either in the MEG Center Tübingen (n = 23) or at Saint Louis University as part of the Human Connectome Project (n = 89) [31]. Two subjects were excluded because of wrong data labeling, and three further subjects were excluded for connectivity analysis because of excessive artifacts. All MEG recordings were performed continuously in magnetically shielded rooms. Fiducials were set to localize the position of the participants’ head in the scanner. EOG and ECG were measured with the MEG.

#### MEG Center data

All participants (n = 23) gave informed consent and the experiment was approved by the local ethics committee. Data was collected during resting state. Participants were asked to sit still and let their mind wander, keeping their eyes open. Each recording lasted 10 minutes. Recordings at the MEG Center were collected with a 275-channel whole-head system (Omega 2000, CTF Systems Inc., Port Coquitlam, Canada). MEG was continuously recorded at a sampling rate of 2343.75 Hz (anti-aliasing filter at Nyquist-frequency).

#### HCP data

The HCP data (S900 release) was from 89 participants (45 female) aged between 22 and 35. All participants gave informed consent and the experiment was approved by the local ethics committee. Resting state data and task data were recorded for each participant. We analyzed the first of the three resting state runs in this study. Each run lasted approximately 6 minutes.

The HCP data was acquired at Saint Louis University using a whole head Magnes 3600 (4D Neuroimaging, San Diego, CA) system having 248 magnetometers and 23 MEG reference channels (5 of them being gradiometers). All measurements were continuously recorded at a sampling rate of 2034.5101 Hz (400 Hz bandwidth, high pass filtered DC) [32].

### Artifact rejection

We adapted the minimally preprocessed pipeline of the HCP [32] to the in-house collected data, and applied the same artifact rejection process to both datasets. The artifact rejection process consisted of three steps. First, we detected bad channels and bad sections of the data. Second, we applied ICA to the data without the bad sections and bad channels and classified each IC as “brain” or artifact using semi-automatic procedures.

Clean data were high-pass filtered at 0.1 Hz using a 4th order forward-reverse Butterworth filter. We removed line noise artifacts with a 1 Hz wide notch filter on 60 or 50 and 51.2 Hz, depending on the provenance of the data, and its harmonic frequencies. Data were finally resampled to 500 Hz.

### Source Projection

We used linearly constrained minimum variance (LCMV) beamforming [48] to project the sensor signals into MNI source space using a single shell model leadfield [49] based on the individual subject’s MRI. The beamforming filter B is obtained using the covariance matrix of the data C of dimensions channels x channels, and the leadfield L of dimensions (3 x channels):

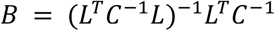

We estimated cortical activity at 457 locations that homogeneously covered the cortical space approximately 7 mm beneath the skull. For each location, we projected the 3-dimensensional source-level data on to its first principal component. We computed the neural activity index (NAI) by dividing the source-level data by a projected noise covariance matrix.

### Broadband signal orthogonalization

IRASA operates on time-domain signals. Thus, to discount signal leakage, we employed pair-wise time-domain orthogonalization of the broadband source-level signals.

We analyzed log-power correlations of time-domain orthogonalized signals to confirm leakage suppression (Fig S2). We split the data into consecutive 3 s windows, orthogonalized each window for each pair of cortical seeds, and computed the power spectral density for each window using FFT. Then, we averaged log transformed power over the following frequency bands: 1−4.5, 4.5−8.75 Hz, 8.75−16 Hz, 16−32 Hz, 32−64 Hz, and 64-125 Hz. We computed the linear correlation of band-limited power between each pair of seeds across time. To test for significant seed patterns of correlation, for each subject and seed, we applied a Fisher z-transform (inverse hyperbolic tangent) and z-scoring to the correlation values to all other seeds. For each seed, we then performed a t-test against 0 across subjects and identified significant clusters with a cluster permutation test.

### IRASA

We applied IRASA to consecutive non-overlapping 3 s windows, which provides an efficient trade-off between computational requirements and spectral resolution [34]. For each window, we performed IRASA using the IRASA software toolbox with default parameters: detrending the data, irregular resampling (parameter *h)* from 1.1 to 1.9 in steps of 0.05, filtering the signal to avoid aliasing when down-sampling, and limiting the output to a frequency range of 0.24 to 125 Hz (one fourth of the sampling rate). For the univariate analysis we performed IRASA per seed. For all pair-wise analyses on the connection level we first orthogonalized the broadband data for each seed pair and then applied IRASA on the orthogonalized signals.

### Modelling the fractal spectrum

For each 3 s window and seed or seed-pair, we fitted a power law to the fractal spectrum obtained from IRASA following the same steps as in the IRASA package. We limited the frequency range from 0.7 to 125 Hz and logarithmically resampled the spectrum. Then we fitted the spectrum with either a one-range linear model, a continuous two-range linear model with a knee at 15 Hz, and a model that takes into account the effect of the ambient noise. For the model including the noise, after fitting the knee model we reversed the logarithm of the predicted model values to add the measured empty room noise multiplied by a noise coefficient *d*. We then again applied the logarithm and computed the mean squared error (MSE) of the model fit. We constrained the noise coefficient between 0 and 3 and minimized the MSE to find the optimal fit. The following function summarizes this model:

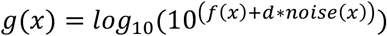

We compared all three models by computing their optimal MSE in the log-linear fitting space and comparing their Akaike information criterion (AIC). The AIC quantifies goodness of fit taking into account the number of model parameters *k* and observations *n*. The AIC was computed as:

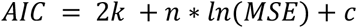

where c was set to 0 for all models.

### Correlation structure of oscillatory and fractal activity

To assess the correlation of different activity components, we log-transformed power spectra and resampled them on a logarithmic frequency axis. The power at frequency f was computed as the inner product between the power spectrum and Gaussian kernels at frequencies f ranging from 1 to 64 Hz in quarter octave steps. The standard deviation of the Gaussian kernel was f/5.83 which results in a spectral bandwidth of 0.5 octaves. Finally, we computed Pearson correlation of power across time for each pair of seeds and frequency. The same approach was used for all signal components, model fits and reconstructed spectra.

### Testing cortical patterns

To quantify whether cortical distributions of model parameters or correlation coefficients were consistent across subjects we performed a cluster permutation statistic (random-effects). First, we z-scored the cortical maps of the model parameters or fisher-z transformed correlation coefficients per subject across space. Then, we performed a t-test against 0 per cortical location across subjects. Clusters were defined as continuous significant regions, with their number of significant locations as the cluster size. Then, 1000 times we randomly flipped the sign of the data of individual subjects and quantified the maximum cluster size for each random permutation. The statistical significance of the original clusters was assessed against the distribution of the permuted maximum cluster sizes.

### Comparing correlation structures

We computed attenuation corrected correlations [8,9,39,40] to compare the correlation structures of fractal and oscillatory components. In short, attenuation correction computes the correlation between those components of two quantities (here: correlations) that are reliable across samples of interest (here: subjects). Critically, attenuation corrected correlations are unbiased, which allows to directly test against 0 and 1.

Frist, we fisher-z transformed and z-scored all cortical correlation matrices. Then we computed the correlation of each seed correlation pattern, i.e. of each column of the correlation matrix, between all pairs of subjects for the fractal and oscillatory components. This yielded the reliabilities rFrac_3,5_ and rOsci_3,5_ where i is a seed and f is a frequency band. We then computed the same correlation of columns between oscillatory and fractal correlation matrices for all pairs of non-identical subjects. This yielded the non-corrected correlation between oscillatory and fractal correlation patterns rOF_3,5_. Finally, we computed the corrected correlation between oscillatory and fractal correlation patterns as 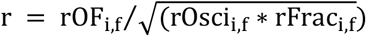. Critically, this correction is only possible for seed patterns, where both oscillatory and fractal component are reliable (i.e. rFrac_3,5_ > 0 and rOsci_3,5_ > 0). Thus, we determined which seeds had reliable correlation patterns using a one-sided t-test across 0. We tested attenuation corrected correlations against 0 and 1 using t-statistics across subjects. Single subject attenuation corrected correlations and reliabilities were estimated as Pseudo-values using a leave-one out Bootstrap.

### Differences between oscillatory and fractal correlation patterns

For each subject and frequency, we vectorized and z-scored the upper triangular matrix of cortical correlation coefficients. Then, for each subject and frequency we subtracted the standardized correlation vector of oscillatory activity from the standardized correlation vector of fractal activity. We then computed attenuation corrected correlations of the resulting difference vectors between frequencies. We computed reliabilities within frequencies across subjects r(*f*_*i*_, *f*_*j*_),we computed the correlation between frequencies r(*f*_*i*_, *f*_*j*_), and we used those values to compute the corrected cross frequency correlation 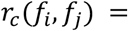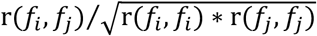.

### Comparison of fractal and oscillatory correlation patterns at the connection level

We computed attenuation corrected correlation between oscillatory and fractal correlation patterns at the connection level [10]. The analysis is analogous to the global attenuation corrected correlation analysis, but, in contrast to the global analysis, which compares the full cortical correlation patterns for each seed, this analysis compares the correlation patterns between the local neighborhoods of each pair of seeds, i.e. of each cortico-cortical connection. Local neighborhoods covered about 19 seed locations within a radius of about 2.5 cm surrounding each seed. This analysis resulted in full cortical correlation matrices of attenuation corrected correlation between oscillatory and fractal correlation patterns. We determined connections with reliabilities > 0 using a t-test on Fisher z-transformed reliabilities (P < 0.01; FDR corrected) and excluded absolute z-scored corrected correlation larger than 4 to stabilize variance.

## Data availability

The HCP data are available for download from https://www.humanconnectome.org/study/hcp-young-adult/document/900-subjects-data-release. MEG Center resting state data is available from the authors upon reasonable request.

## Acknowledgements

We thank Marcus Siems and Janet Giehl for helpful discussions and insights. This work was supported by the European Research Council (ERC) StG335880 (M.S) and the Centre for Integrative Neuroscience (DFG, EXC 307) (M.S.). The authors acknowledge support by the state of Baden-Württemberg through bwHPC and the German Research Foundation (DFG) through grant no INST 39/963-1 FUGG (bwForCluster NEMO).

## Author Contributions

Conceptualization: M.S., A.I.C; Methodology: A.I.C; Investigation: A.I.C.; Formal Analysis, A.I.C; Writing – Original Draft: A.I.C, M.S.; Writing – Review & Editing: A.I.C, M.S.; Funding Acquisition: M.S.; Resources: M.S.; Supervision, M.S.

## Supplementary figures

**Fig S1.**
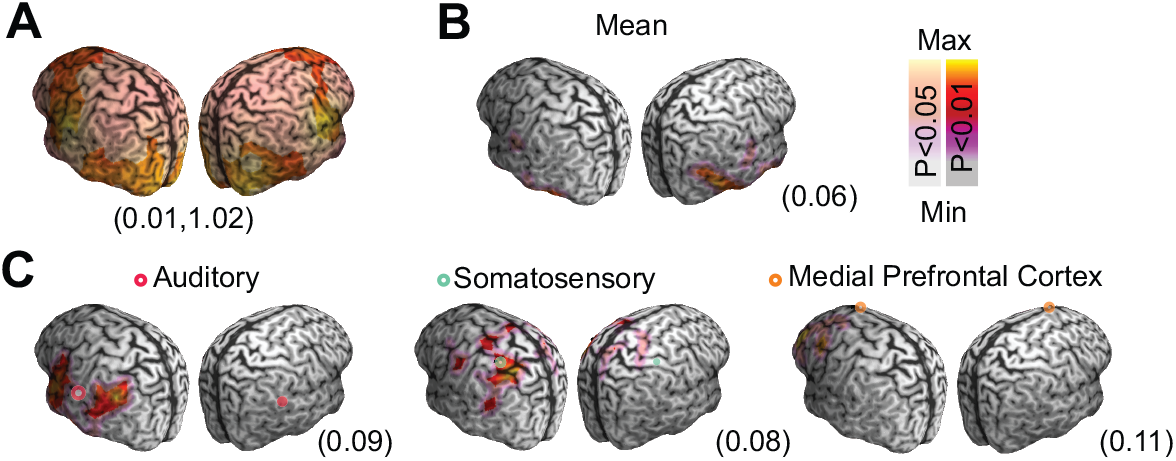
Noise component of the fractal power model. (**A**) Distribution of the noise coefficient for the fractal model over the cortex. Values in parentheses indicate minimum and maximum. Hight noise coefficient values concentrate over temporal cortex, areas prone to residual muscle artifacts. (**B**) Average cortical correlation of the noise coefficient marginalized across seeds. Value in parentheses indicate maximum. (**C**) Seed-correlation patterns of the noise component for three characteristic seeds. Maxima in parentheses. All panels show cluster permutation statistics.

**Fig S2.**
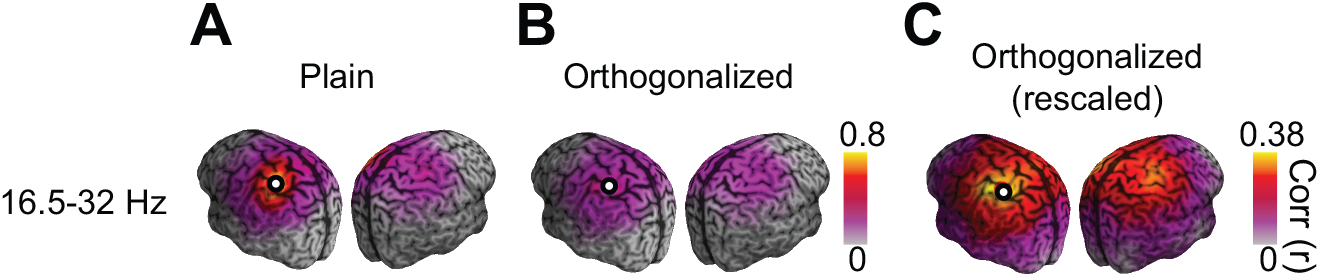
Time-domain orthogonalization discounts spurious coupling induced by signal leakage. (**A**) Correlation pattern of 16-32 Hz activity for left somatosensory cortex without orthogonalization. (**B**) Correlation with time-domain orthogonalization. (**C**) Same result as in panel (A) but colors rescaled to min max.

**Fig S3.**
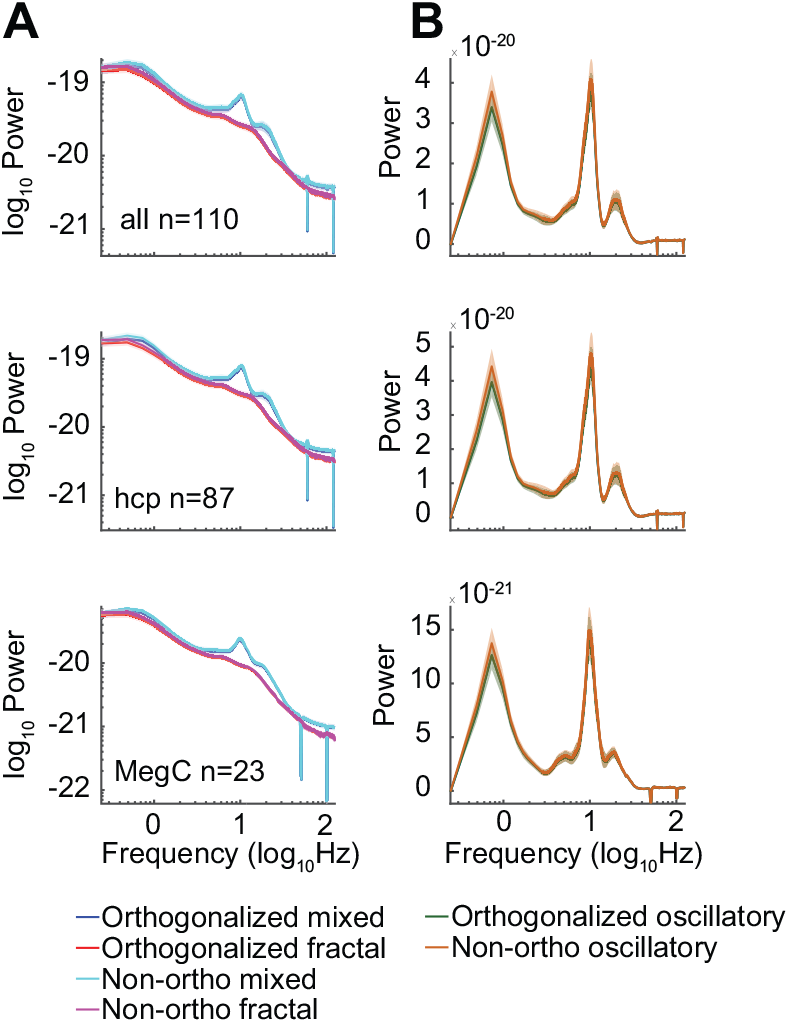
Orthogonalization does not alter the power spectrum shape. (**A**) Marginal mean of orthogonalized mixed and fractal power compared to non-orthogonalized power distributions, mean over space and subjects. From top to bottom: all subjects, HCP subjects, Meg Center subjects. (**B**) Same as in (A) but for oscillatory power. Shaded areas denote SEM across subjects.

**Fig S4.**
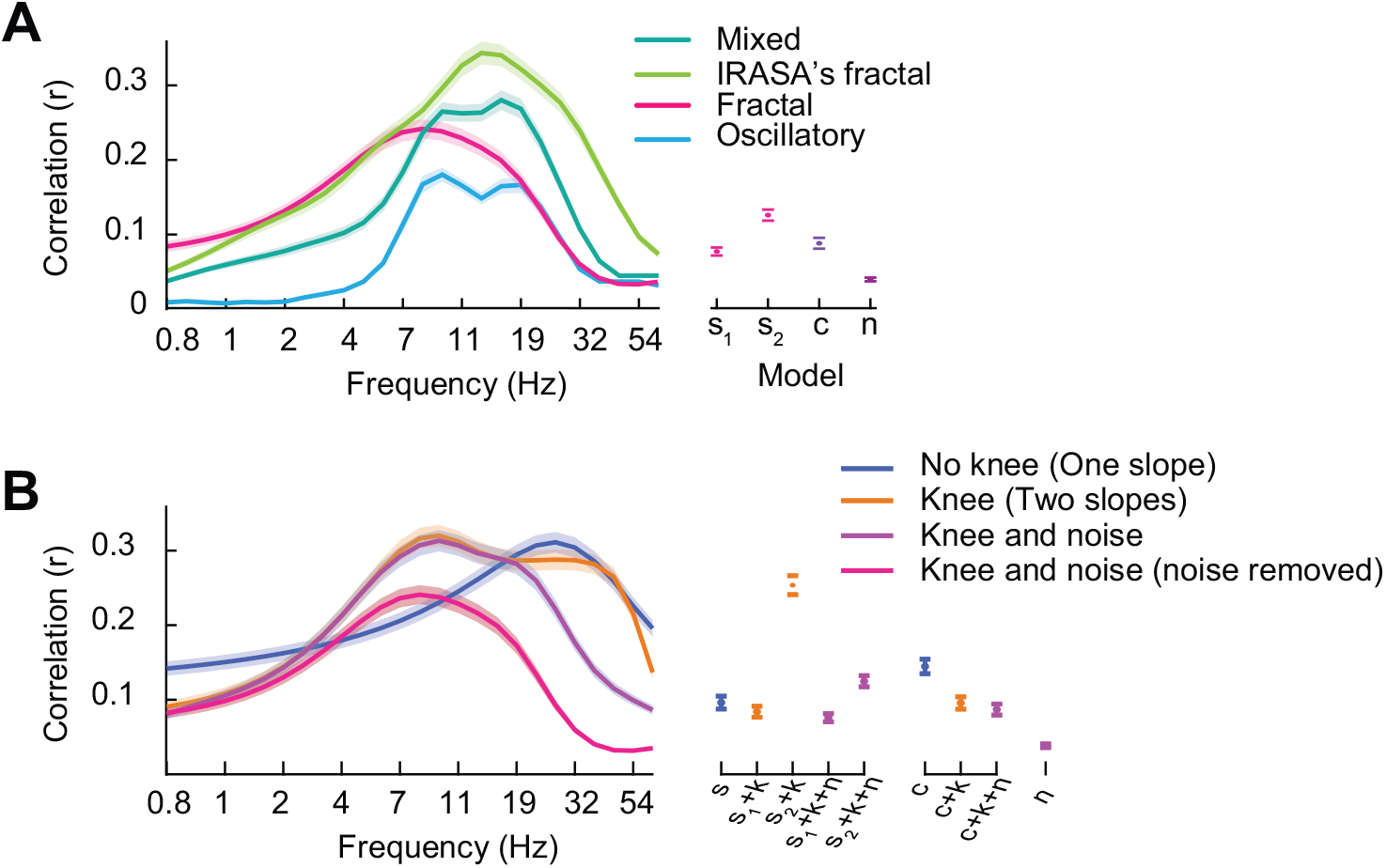
Spectral profile of cortical correlations of different fractal models. (**A**) Mean cortical correlation for the oscillatory, mixed, IRASA’s fractal and the fractal model. On the right are the correlations for the four coefficients of the fractal model. Same data as in Fig 4A. (**B**) Mean cortical correlation of the different fractal models and their corresponding parameters. Shaded areas and error bars denote SEM across subjects.

**Fig S5.**
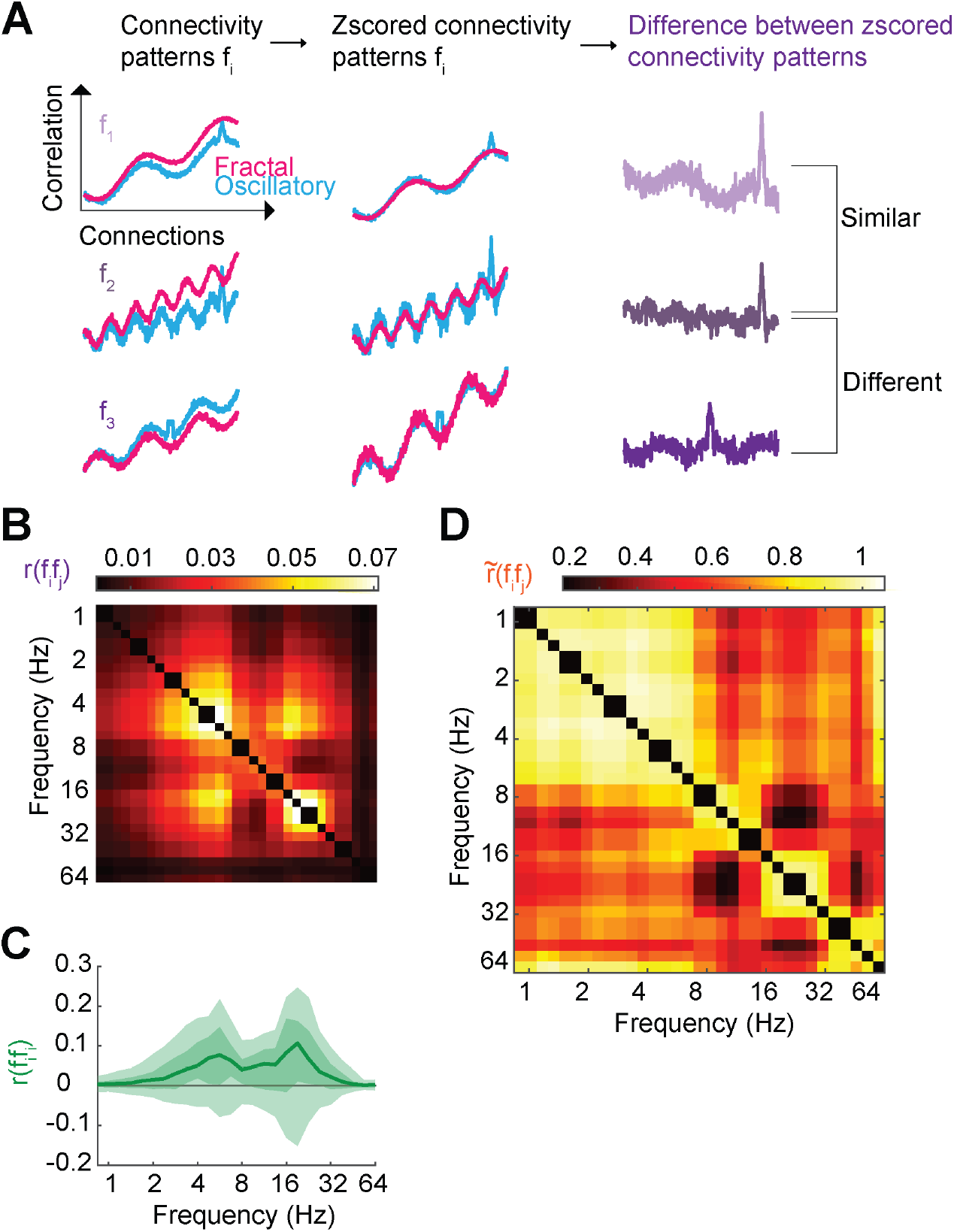
Comparison of differences of correlation patterns across frequencies. (**A**) Schematic illustration. Subtracting z-scored correlation patterns reveals local distinctions beyond otherwise similar global correlations structures. (**B**) Uncorrected correlation of differences (fractal-oscillatory) of correlation patterns across frequencies. (**C**) Reliabilities of differences across subjects within each frequency. Shaded areas denote 5-95^th^ and 25-75^th^ percentiles. (**D**) Attenuation corrected correlation of differences across frequencies (same as Fig 6B).

**Fig S6.**
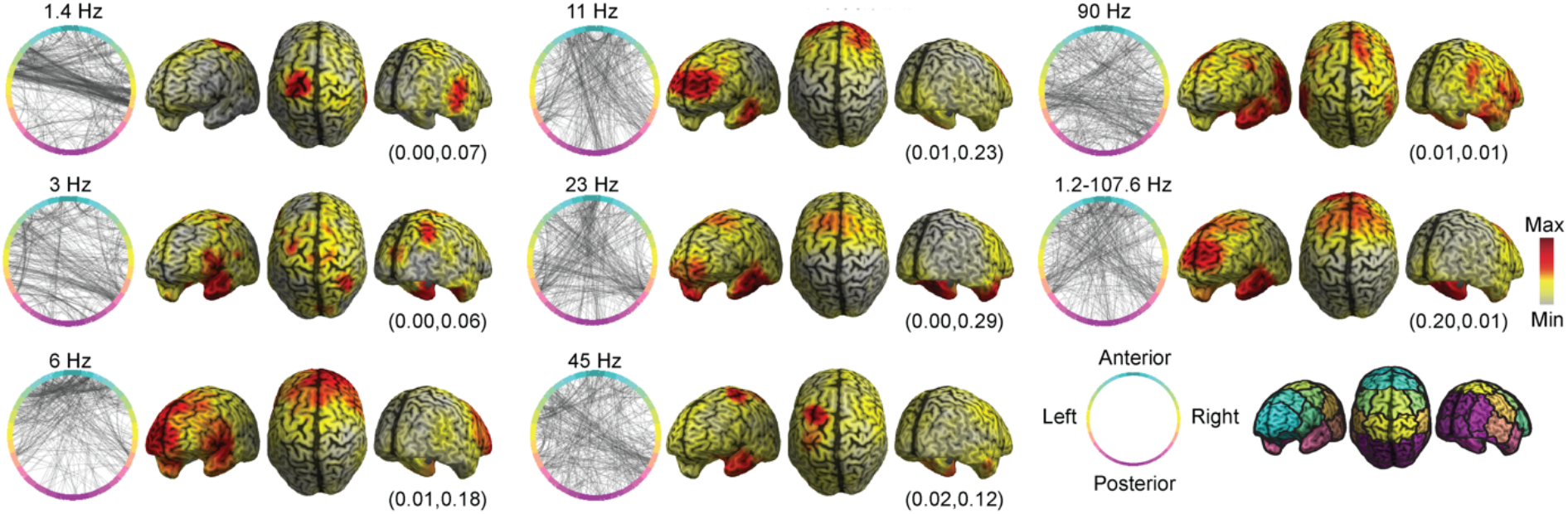
Local differences between fractal and oscillatory correlations. Circle plots show a random sample (800 connections) of the 10 % lowest average corrected correlations across frequency-bands and subjects. The topographies show the marginal mean of the corrected correlations that are in the 10 % lowest correlations per frequency band. Minimum and maximum marginal correlations are in parentheses. The average across the full frequency range is shown on the middle right.

